# PD-L1 expression in equine malignant melanoma and functional effects of PD-L1 blockade

**DOI:** 10.1101/2020.05.22.110395

**Authors:** Otgontuya Ganbaatar, Satoru Konnai, Tomohiro Okagawa, Yutaro Nojima, Naoya Maekawa, Erina Minato, Atsushi Kobayashi, Ryo Ando, Nobuya Sasaki, Daisuke Miyakosi, Osamu Ichi, Yukinari Kato, Yasuhiko Suzuki, Shiro Murata, Kazuhiko Ohashi

## Abstract

Programmed death-1 (PD-1) is an immunoinhibitory receptor expressed on exhausted T cells during chronic illness. Interaction of PD-1 with its ligand PD-ligand 1 (PD-L1) delivers inhibitory signals and impairs proliferation, cytokine production, and cytotoxicity of T cells. We reported that the PD-1/PD-L1 pathway is closely associated with T-cell exhaustion and disease progression in bovine chronic infections and canine tumors. Moreover, we found that blocking antibodies targeting PD-1 and PD-L1 restore T-cell functions and may be used in immunotherapy in cattle and dogs. However, the immunological role of the PD-1/PD-L1 pathway remains unclear for chronic equine diseases, including tumors. In this study, we identified nucleotide sequences of equine PD-1 (EqPD-1) and PD-L1 (EqPD-L1) and investigated the role of anti-bovine PD-L1 monoclonal antibodies (mAbs) against EqPD-L1 using *in vitro* assays. We also evaluated the expression of PD-L1 in tumor tissues of equine malignant melanoma (EMM).

The amino acid sequences of EqPD-1 and EqPD-L1 share a high identity and similarity with homologs from other mammalian species. Two clones of the anti-bovine PD-L1 mAbs recognized EqPD-L1 in flow cytometry, and one of these cross-reactive mAbs blocked the binding of equine PD-1/PD-L1. Importantly, PD-L1 expression was confirmed in EMM tumor tissues by immunohistochemistry. A cultivation assay revealed that PD-L1 blockade enhanced the production of Th1 cytokines in equine immune cells.

These results suggest that our anti-PD-L1 mAbs may be useful for investigating the expression and role of the equine PD-1/PD-L1 pathway. Further research is required to discover the immunological role of PD-1/PD-L1 in chronic equine diseases and elucidate a future application in immunotherapy for horse.

## Introduction

Programmed cell death-1 (PD-1) is an immunoinhibitory receptor, which is expressed on activated and exhausted T cells [1]. Its ligand programmed death ligand 1 (PD-L1) is expressed on immune cells, including antigen-presenting cells, and tumor cells [1]. The interaction of PD-1 and PD-L1 suppresses the activation signal mediated by T-cell receptors and inhibits effector functions of T cells, such as cytokine production and cell proliferation [1]. This pathway is invaluable for regulating excessive immune responses. In cancers, however, tumor cells utilize the suppression of T cells mediated by PD-1/PD-L1 to avoid anti-tumor immune responses [1]. In human medicine, the blocking antibodies targeting PD-1 or PD-L1 have been leveraged for treatment of various types of cancers and resulted in remarkable outcomes with 20%–90% response rates in multiple clinical trials [1].

Equine malignant melanoma (EMM) is a common neoplasm among aged gray horses, which results in dermal tumors at multiple sites [2]. A previous study reported that around 80% of aged gray horses developed dermal melanoma and predicted that all gray horses would develop this tumor as they reach old age [3]. Cellular immune response is critical for the eradication of melanoma, but several mechanisms have been propounded to limit anti-tumor immunity in EMM based on the findings for human malignant melanoma [4]. However, no studies on immune evasion mechanisms in EMM have yet been done, and immune exhaustion mediated by PD-1 and PD-L1 has not been investigated in horses.

In our previous research, we established anti-bovine PD-L1 monoclonal antibodies (mAbs) [5]. We found that PD-1 and PD-L1 play critical roles in immune exhaustion and disease progression in bovine chronic infections [6–10] and in canine cancers including malignant melanoma [11, 12]. Importantly, we noted that the PD-1/PD-L1 blockade enhances T-cell responses in cattle and dogs and exhibits therapeutic effects in bovine chronic infections and canine malignant melanoma [6–10, 13–17].

So far, there are no reports on genetic information, expression, and function of PD-1/PD-L1 in horses. Furthermore, the role of the PD-1/PD-L1 pathway in EMM remains unclear. Based on the findings of our previous studies, we hypothesized that PD-1 and PD-L1 may provide potential targets for immunotherapy against EMM. Hence, in this study, we identified nucleotide sequences of equine PD-1 (EqPD-1) and PD-L1 (EqPD-L1), evaluated the blocking effects of our anti-bovine PD-L1 mAbs against EqPD-L1, and confirmed the expression of PD-L1 on EMM.

## Materials and Methods

### Horse blood samples and cell preparation

Heparinized blood samples were collected from Thoroughbred horses in farms and veterinary hospitals in Hokkaido, Japan. Peripheral blood mononuclear cells (PBMCs) were purified using density gradient centrifugation on Percoll (GE Healthcare, Little Chalfont, UK), washed three times with phosphate-buffered saline (PBS), and suspended in PBS. All experimental procedures were conducted following approval from the local committee for animal studies according to the Hokkaido University (17-0024). Informed consent was obtained from all owners.

### Cloning of cDNA encoding of equine PD-1 and PD-L1

Equine PBMCs (4 × 10^6^ cells) were cultivated with 20 ng/mL of phorbol 12-myristate acetate (PMA; Sigma–Aldrich, St. Louis, MO, USA) and 1 μg/mL of ionomycin (Sigma–Aldrich) in RPMI 1640 medium (Sigma–Aldrich) supplemented with 10% heat-inactivated fetal bovine serum (FBS) (Thermo Fisher Scientific, Waltham, MA, USA), 2 mM of L-glutamine, 100 U/mL of penicillin, and 100 μg/mL of streptomycin (Thermo Fisher Scientific) at 37°C with 5% CO_2_ for 24 h.

Total RNA was isolated from cultivated PBMCs using of TRI Reagent (Molecular Research Center, Cincinnati, OH, USA) according to the manufacturer’s instructions. Residual DNA was removed from RNA samples by treatment with Deoxyribonuclease I (Thermo Fisher Scientific). cDNA was synthesized from 1 μg of total RNA with PrimeScript Reverse Transcriptase (Takara Bio, Otsu, Japan) and oligo(dT) primer according to the manufacturer’s instructions.

Gene-specific primers were designed to amplify EqPD-1 and EqPD-L1 genes, based on the sequences from horses available on GenBank (XM_005610777 and XM_001492842). EqPD-1 and EqPD-L1 cDNAs were amplified by PCR using TaKaRa Ex Taq (Takara Bio) and specific primers (Supplementary Table 1). The PCR products were purified using a FastGene Gel/PCR Extraction Kit (Nippon Genetics, Tokyo, Japan), and cloned into the pGEM-T Easy Vector (Promega, Madison, WI, USA). They were transferred into *E. coli* HST08 Premium Competent Cells (Takara Bio) and plated onto LB agar plates (Sigma–Aldrich) containing X-gal (Takara Bio) and ampicillin (Sigma–Aldrich). The purified plasmid clones were sequenced using a GenomeLab GeXP Genetic Analysis System (SCIEX, Framingham, MA, USA). The established sequences were aligned, and an unrooted neighbor-joining tree was constructed using MEGA software program version 7.0 [17].

### Preparation of EqPD-1- and EqPD-L1-expressing cells

cDNAs encoding EqPD-1 and EqPD-L1 were amplified by PCR using gene-specific primers with restriction enzyme cleavage sites (Supplementary Table 1) and subcloned into the multicloning site of pEGFP-N2 (Clontech, Palo Alto, CA, USA).

COS-7 cells were cultured in RPMI 1640 medium (Sigma–Aldrich) supplemented with 10% heat-inactivated FBS (Thermo Fisher Scientific), 2 mM of L-glutamine, 100 U/mL of penicillin, and 100 μg/mL of streptomycin (Thermo Fisher Scientific) at 37°C and 5% CO_2_. The cells were transfected with purified plasmids using Lipofectamine 3000 Reagent (Thermo Fisher Scientific) and cultivated for 48 h after transfection. The cellular localization of EqPD-1-EGFP and EqPD-L1-EGFP was then confirmed using the ZOE Fluorescent Cell Imager (Bio-Rad, Hercules, CA, USA).

### Expression and purification of soluble equine PD-1 and PD-L1 proteins

Soluble forms of EqPD-1 and EqPD-L1 proteins fused with rabbit IgG Fc region (EqPD-1-Ig and EqPD-L1-Ig) were obtained by amplifying cDNAs encoding the extracellular domain fragments of EqPD-1 and EqPD-L1 with signal sequences by PCR with gene-specific primers with restriction enzyme cleavage sites (Supplementary Table 1). The amplicons were subcloned into the multicloning site of pCXN2.1(+) (kindly provided by Dr. T. Yokomizo, Juntendo University, Japan) [19] with the gene cassette encoding the Fc region of rabbit IgG. Transient cell lines expressing EqPD-1-Ig and EqPD-L1-Ig were established with the use of an Expi293 Expression System (Thermo Fisher Scientific). Expi293F cells were transfected with pCXN2.1(+)-EqPD-1-Ig and pCXN2.1(+)-EqPD-L1-Ig using Expifectamine (Thermo Fisher Scientific) and cultivated with shaking in Expi293 medium (Thermo Fisher Scientific) at 37°C and 125 rpm with 8% CO_2_ for 7 days.

Purification of EqPD-1-Ig and EqPD-L1-Ig from the culture supernatants was achieved by affinity chromatography with an Ab-Capcher ExTra (ProteNova, Kagawa, Japan). The buffer was exchanged with PBS by size exclusion chromatography using a PD-10 Desalting Column (GE Healthcare). The purity of EqPD-1-Ig and EqPD-L1-Ig was confirmed by sodium dodecyl sulfate-polyacrylamide gel electrophoresis (SDS-PAGE) in reducing or nonreducing conditions using SuperSep Ace 5%–20% gradient polyacrylamide gel (FUJIFILM Wako Pure Chemical, Osaka, Japan) and 2 × Laemmli Sample Buffer (Bio-Rad).

Precision Plus Protein All Blue Standard (Bio-Rad) was used as a molecular-weight size marker. The proteins were visualized with Quick-CBB (FUJIFILM Wako Pure Chemical), and protein concentrations were measured by ultraviolet absorbance at 280 nm with a NanoDrop 8000 Spectrophotometer (Thermo Fisher Scientific).

### Binding assay of EqPD-1 and EqPD-L1

Binding of EqPD-1-Ig and EqPD-L1-Ig to COS-7 cells expressing EqPD-L1-EGFP and EqPD-1-EGFP were investigated using flow cytometry. EqPD-L1-EGFP cells or EqPD-1-EGFP cells were incubated with 10 μg/mL of biotinylated EqPD-1-Ig or EqPD-L1-Ig, respectively, at 37°C for 30 min. Biotinylated rabbit control IgG (Southern Biotech, Birmingham, AL, USA) was used as a negative control. EqPD-1-Ig, EqPD-L1-Ig, and rabbit control IgG were biotinylated using a Lightning-Link Rapid Type A Biotin Conjugation Kit (Innova Biosciences, Cambridge, UK). Cells were washed with PBS containing 1% bovine serum albumin (BSA; Sigma–Aldrich) and labeled using APC-conjugated streptavidin (BioLegend, San Diego, CA, USA) at 25°C for 30 min. After rewashing, cells were immediately analyzed by FACS Verse (BD Biosciences, San Jose, CA, USA).

### Cross-reactivity assay of anti-bovine PD-L1 mAbs against EqPD-L1

EqPD-L1-EGFP cells were incubated with four clones of anti-bovine PD-L1 mAbs (4G12-C1, rat IgG_2a_; 5A2-A1, rat IgG_1_; 6C11-3A11, rat IgG_2a_; 6G7-E1, rat IgM) [5, 20] at 25°C for 20 min to analyze the binding ability of anti-bovine PD-L1 mAbs to EqPD-L1. Rat IgG_1_ (R3-34, BD Biosciences, San Jose, CA, USA), rat IgG_2a_ (R35-95, BD Biosciences), and rat IgM isotype controls (R4-22, BD Biosciences) were used for negative control staining. Cells were washed with 1% BSA-PBS and labeled with APC-conjugated goat anti-rat immunoglobulin antibody (Southern Biotech) at 25°C for 20 min. After rewashing, cells were immediately analyzed by FACS Verse (BD Biosciences).

Fresh and stimulated equine PBMCs were analyzed by flow cytometry to analyze the binding ability of anti-bovine PD-L1 mAbs to equine immune cells. Equine PBMCs (4 × 10^6^ cells) were stimulated in cultivation with 20 ng/mL of PMA (Sigma–Aldrich) and 1 μg/mL of ionomycin (Sigma–Aldrich) for 24 h, as described above. Fresh and stimulated PBMCs were incubated with PBS supplemented with 10% goat serum (Thermo Fisher Scientific) at room temperature for 15 min to prevent nonspecific reactions. Cells were washed and stained with anti-PD-L1 mAbs (5A2-A1, rat IgG_1_; 6C11-3A11; rat IgG_2a_) [5, 20] at room temperature for 30 min. Rat IgG_1_ (R3-34, BD Biosciences) and rat IgG_2a_ isotype controls (R35-95, BD Biosciences) were used for negative control staining. Cells were washed with PBS containing 1% BSA (Sigma–Aldrich) and labeled with APC-conjugated anti-rat Ig antibody (Southern Biotech) at room temperature for 30 min. After rewashing, cells were immediately analyzed by FACS Verse (BD Biosciences).

### Blockade assay of EqPD-1/EqPD-L1 interaction

Blocking assays were conducted on microplates using EqPD-1-Ig and EqPD-L1-Ig to analyze the ability of the anti-PD-L1 mAbs to block PD-1/PD-L1 binding. MaxiSorp Immuno Plates (Thermo Fisher Scientific) were coated with EqPD-1-Ig (1 μg/mL) in carbonate-bicarbonate buffer (Sigma–Aldrich) and blocked with SuperBlock T20 (PBS) Blocking Buffer (Thermo Fisher Scientific). Biotinylated EqPD-L1-Ig was preincubated with anti-PD-L1 mAb 5A2-A1 (rat IgG_1_) [5], 6C11-3A11 (rat IgG_2a_) [20], rat IgG_1_ isotype control (R3-34, BD Biosciences), or rat IgG_2a_ isotype control (R35-95, BD Biosciences) at various concentrations (0, 1.25, 2.5, 5.0, 7.5, 10 μg/mL) at 37°C for 30 min. The preincubated reagents were added to the microplates and incubated at 37°C for a further 30 min. EqPD-L1-Ig binding was detected using horseradish peroxidase-conjugated Neutravidin (Thermo Fisher Scientific) and TMB One Component Substrate (Bethyl Laboratories, Montgomery, TX, USA). Optical density at 450 nm was measured by a microplate reader MTP-900 (Corona Electric, Hitachinaka, Japan). Three independent experiments were each performed in duplicate.

### Immunohistochemical assay of PD-L1

Tumor tissues from four horses bearing EMM were immunohistochemically stained (Supplementary Table 2). The tissues were fixed in formalin, embedded into paraffin wax and cut into 4-μm-thick sections. The dried sections were deparaffinized in xylene and hydrated through graded alcohols. Melanin was bleached from the sections with using 0.25% potassium permanganate and 2% oxalic acid. Antigen retrieval was achieved using 0.01 M citrate buffer (pH 6.0) by microwave heating. Endogenous peroxidase activity was blocked by incubating the sections in methanol containing 0.3% hydrogen peroxide. The sections were incubated with or without anti-PD-L1 mAb (6C11-3A11, rat IgG_2a_) [20] at 4°C overnight, followed by detection using Vectastain Elite ABC Rat IgG kit (Vector Laboratories, Burlingame, CA, USA). The immunoreaction was visualized using 3,3’-diaminobenzidine tetrahydrochloride. All immunostained sections were examined under an optical microscope.

### Immunoactivation assay using equine PBMCs

To determine the effects of inhibiting the PD-1/PD-L1 interaction on equine immune cells, equine PBMCs were cultured with 10 μg/mL of anti-PD-L1 mAb (6C11-3A11, rat IgG_2a_) [20] or rat IgG_2a_ control (Bio X Cell, West Lebanon, NH, USA) in the presence of 0.1 μg/mL of Staphylococcal enterotoxin B (Sigma–Aldrich) at 37°C with 5% CO_2_ for three days. All cell cultures were grown in 96-well round-bottomed plates (Corning Inc., Corning, NY, USA) containing 4 × 10^5^ PBMCs in 200 μl of RPMI 1640 medium (Sigma–Aldrich) supplemented with 10% heat-inactivated FBS, 2 mM of L-glutamine, 100 U/mL of penicillin, and 100 μg/mL of streptomycin (Thermo Fisher Scientific). Cytokine concentrations in the culture supernatants were determined using an Equine IFN-γ ELISA Development Kit (Mabtech, Nacka Strand, Sweden) and an Equine IL-2 DuoSet ELISA (R&D Systems, Minneapolis, MN, USA). Measurements were performed in duplicate according to the manufacturer’s protocol.

### Statistical analysis

Significant differences were identified using Wilcoxon signed-rank test or Tukey’s test. All statistical tests were performed using the MEPHAS (http://www.gen-info.osaka-u.ac.jp/MEPHAS/) statistical analysis program. Statistical significance was set as *p* < 0.05.

## Results

### Molecular cloning and sequence analysis of equine PD-1/PD-L1

Figs 1A and 2A show the putative amino acid sequences of EqPD-1 and EqPD-L1, respectively. EqPD-1 and EqPD-L1 consist of a putative signal peptide, an extracellular region, a transmembrane region, and an intracellular region. These were expected to be type I transmembrane proteins as orthologues in other species. A conserved domain search identified an immunoglobulin variable (IgV)-like domain in the extracellular regions of EqPD-1. IgV-like and immunoglobulin constant (IgC)-like domains were observed in the extracellular regions of EqPD-L1. The intracellular region of EqPD-1 contained two structural motifs, an immunoreceptor tyrosine-based inhibitory motif (ITIM) and an immunoreceptor tyrosine-based switch motif (ITSM). Phylogenetic analyses revealed that EqPD-1 and EqPD-L1 were clustered in a group comprising Artiodactyla and Carnivora (Figs 1B and 2B). EqPD-1 had 70.4%, 69.0%, 75.6%, 69.2%, and 58.7% amino acid similarities to pig, cattle, dog, human, and mouse respectively (Table 1). EqPD-L1 amino acid similarities to pig, cattle, dog, human, and mouse were 81.5%, 80.9%, 83.7%, 79.0%, and 67.9%, respectively (Table 2).

**Table 1.**
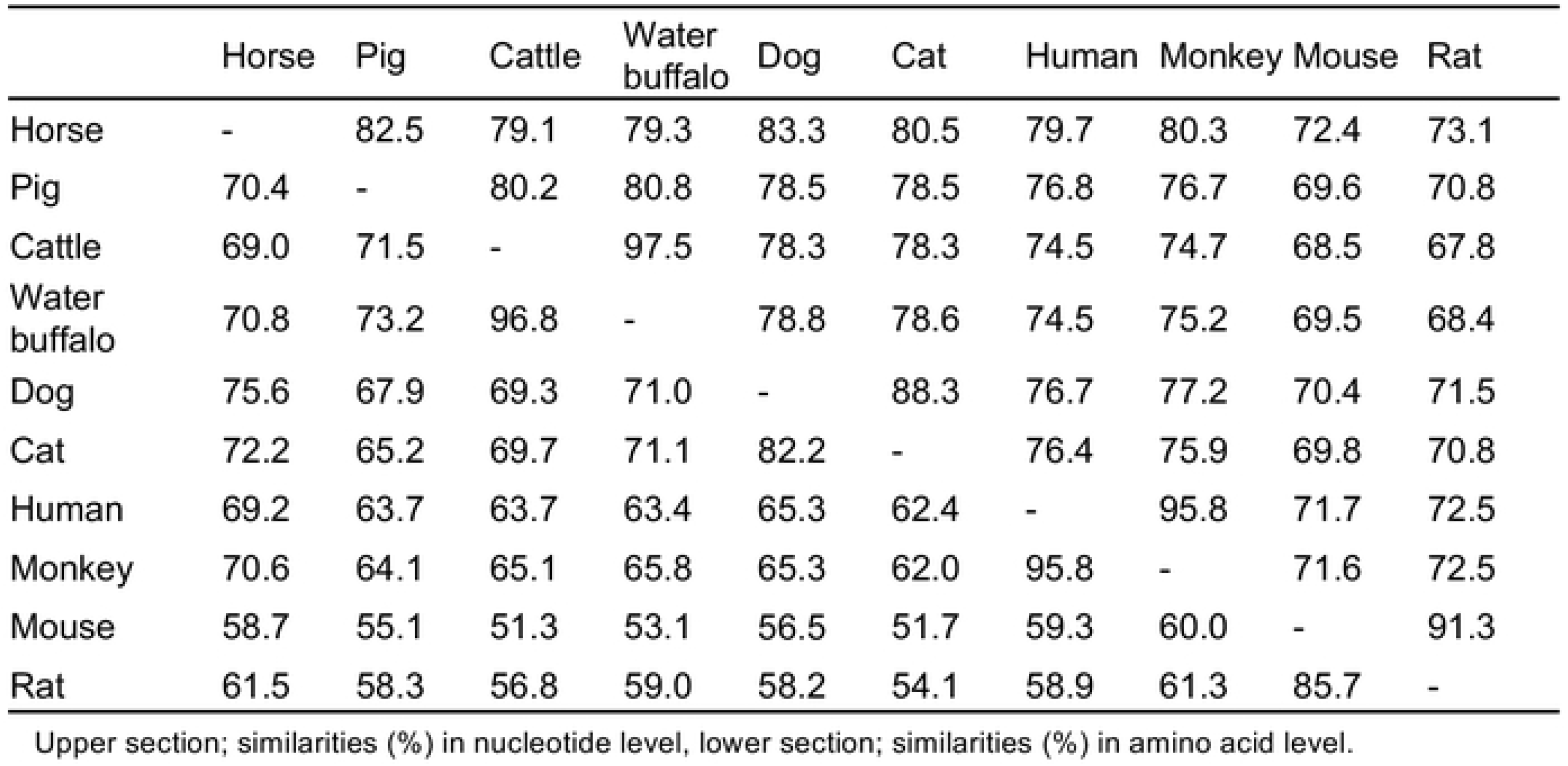
Similarities of nucleotide and amino acid sequences of PD-1 among mammalian species.

|  | Horse | Pig | Cattle | Water buffalo | Dog | Cat | Human | Monkey | Mouse | Rat |
| --- | --- | --- | --- | --- | --- | --- | --- | --- | --- | --- |
| Horse | - | 82.5 | 79.1 | 79.3 | 83.3 | 80.5 | 79.7 | 80.3 | 72.4 | 73.1 |
| Pig | 70.4 | - | 80.2 | 80.8 | 78.5 | 78.5 | 76.8 | 76.7 | 69.6 | 70.8 |
| Cattle | 69.0 | 71.5 | - | 97.5 | 78.3 | 78.3 | 74.5 | 74.7 | 68.5 | 67.8 |
| Water buffalo | 70.8 | 73.2 | 96.8 | - | 78.8 | 78.6 | 74.5 | 75.2 | 69.5 | 68.4 |
| Dog | 75.6 | 67.9 | 69.3 | 71.0 | - | 88.3 | 76.7 | 77.2 | 70.4 | 71.5 |
| Cat | 72.2 | 65.2 | 69.7 | 71.1 | 82.2 | - | 76.4 | 75.9 | 69.8 | 70.8 |
| Human | 69.2 | 63.7 | 63.7 | 63.4 | 65.3 | 62.4 | - | 95.8 | 71.7 | 72.5 |
| Monkey | 70.6 | 64.1 | 65.1 | 65.8 | 65.3 | 62.0 | 95.8 | - | 71.6 | 72.5 |
| Mouse | 58.7 | 55.1 | 51.3 | 53.1 | 56.5 | 51.7 | 59.3 | 60.0 | - | 91.3 |
| Rat | 61.5 | 58.3 | 56.8 | 59.0 | 58.2 | 54.1 | 58.9 | 61.3 | 85.7 | - |

Upper section; similarities (%) in nucleotide level, lower section; similarities (%) in amino acid level.

**Table 2.** Similarities of nucleotide and amino acid sequenc es of PD-L1 among mammalian spec ies.

|  | Horse | Pig | Cattle | Water buffalo | Dog | Human | Monkey | Mouse | Rat |
| --- | --- | --- | --- | --- | --- | --- | --- | --- | --- |
| Horse | - | 88.8 | 87.8 | 87.3 | 88.1 | 86.6 | 85.2 | 75.6 | 75.7 |
| Pig | 81.5 | - | 88.1 | 88.2 | 85.1 | 84.3 | 83.3 | 74.4 | 75.5 |
| Cattle | 80.9 | 81.3 | - | 98.2 | 84.0 | 83.0 | 81.8 | 73.8 | 75.2 |
| Water buffalo | 79.9 | 80.6 | 96.5 | - | 83.7 | 82.3 | 81.5 | 74.2 | 75.7 |
| Dog | 83.7 | 76.8 | 78.5 | 77.8 | - | 83.2 | 82.1 | 73.4 | 74.5 |
| Human | 79.0 | 73.3 | 73.1 | 72.0 | 75.9 | - | 95.7 | 76.3 | 76.4 |
| Monkey | 76.6 | 71.6 | 72.7 | 72.0 | 74.2 | 91.3 | - | 75.2 | 75.3 |
| Mouse | 67.9 | 67.4 | 65.1 | 66.2 | 67.0 | 69.4 | 68.3 | - | 87.3 |
| Rat | 68.3 | 67.4 | 67.3 | 67.6 | 68.1 | 69.7 | 68.3 | 83.4 | - |

Upper section; similarities (%) in nucleotide level, lower section; similarities (%) in amino acid level.

**Figure 1.**
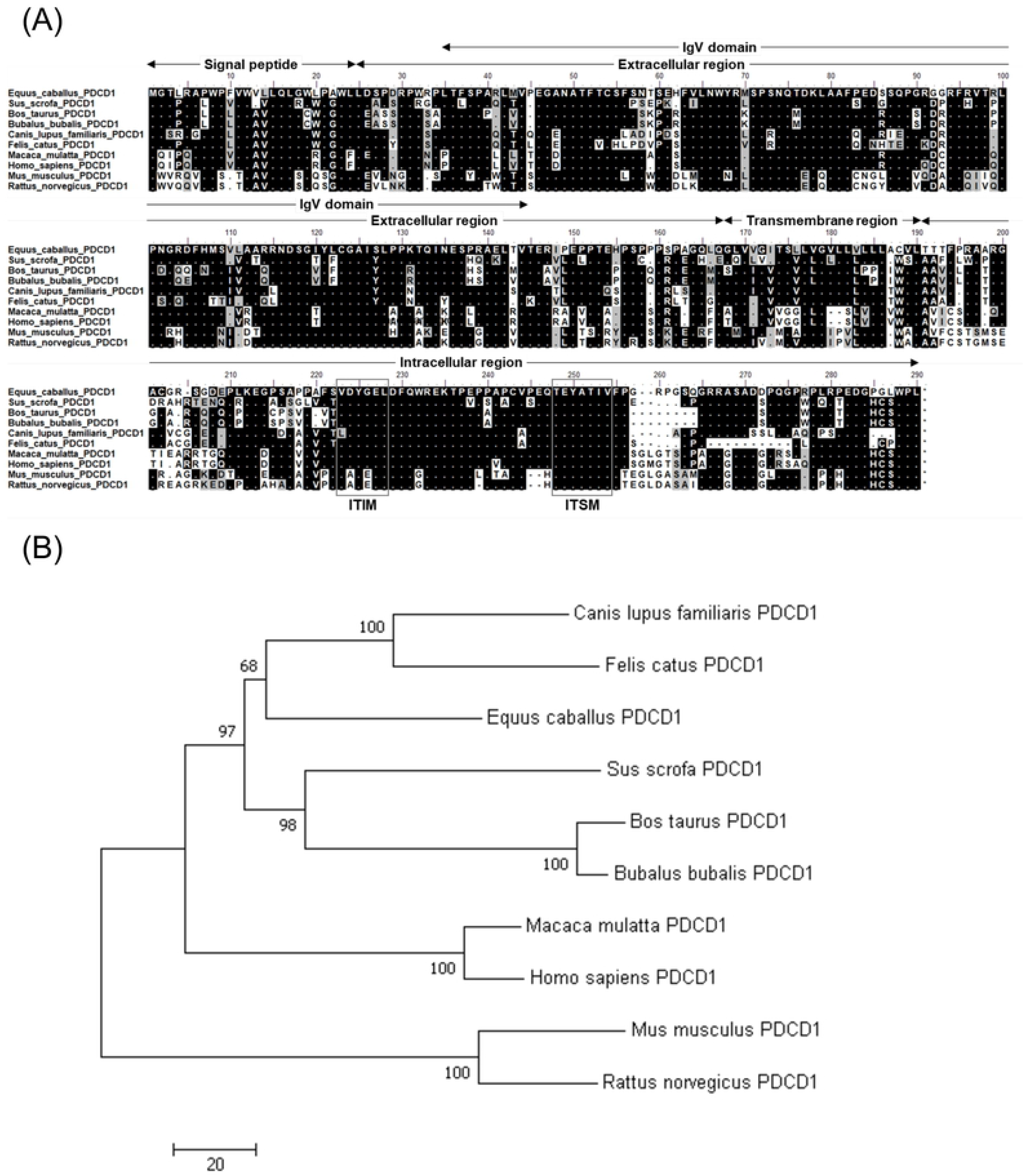
Sequence analysis of EqPD-1. (A) Multiple sequence alignment of amino acid sequences of equine and vertebrate PD-1. Predicted domains and motifs of EqPD-1 are shown. EqPD-1 consists of a signal peptide, an extracellular region, a transmembrane region, and an intracellular region. The cytoplasmic tail of PD-1 contains the ITIM and ITSM motifs. (B) Phylogenetic tree of EqPD-1 sequence in relation to those of other vertebrate species. The bootstrap consensus tree was inferred from 1000 replicates with the neighbor-joining method using the MEGA 7.0 software. The scale indicates the divergence time. The GenBank accession numbers of nucleotide sequences used in these analyses are listed in Supplementary Table 3.

**Figure 2.**
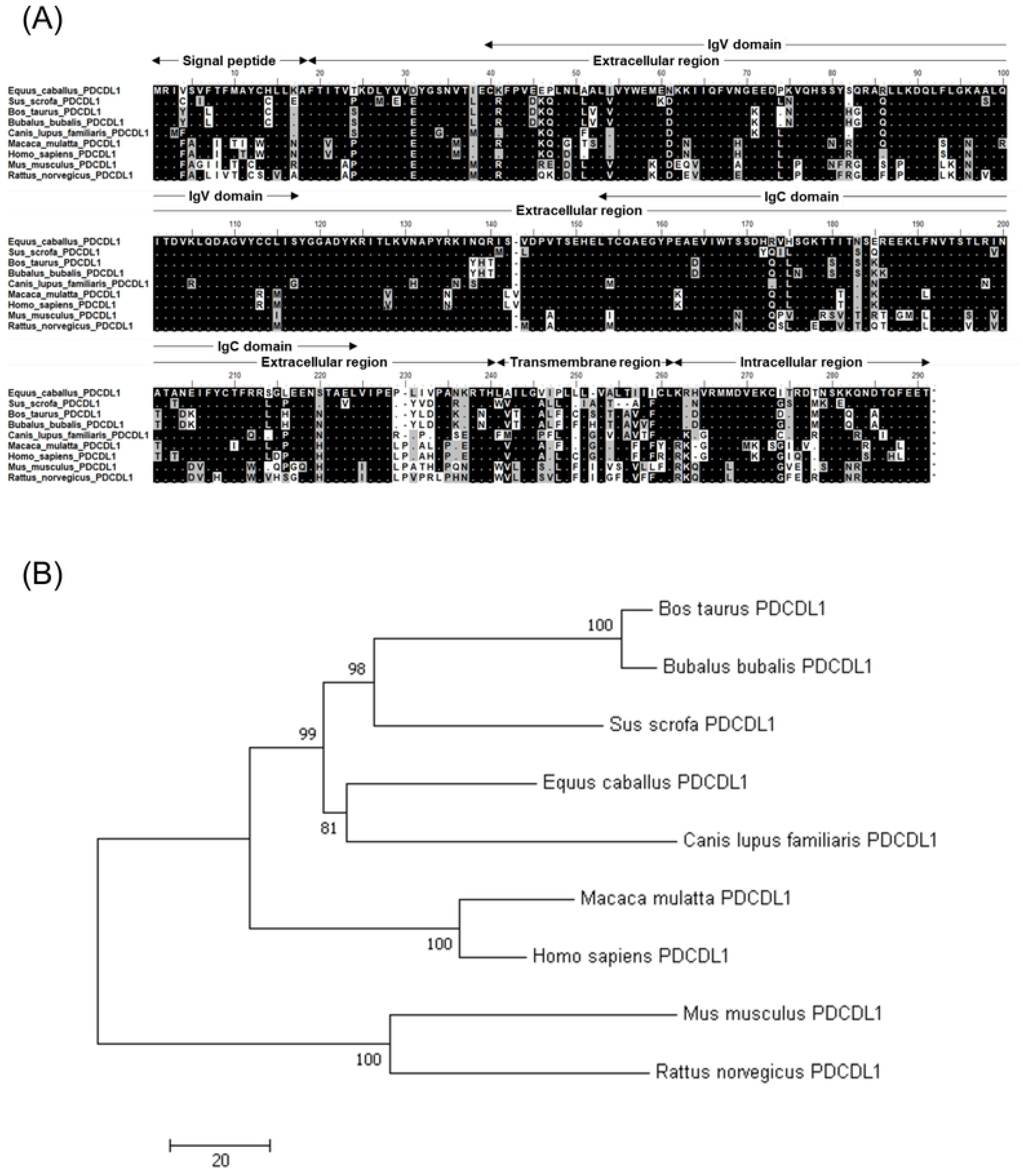
Sequence analysis of EqPD-L1. (A) Multiple sequence alignment of PD-L1 amino acid sequences of equine and vertebrate PD-1. Predicted domains and motifs of EqPD-L1 are shown in the figure. EqPD-L1 consists of a signal peptide, an extracellular region, a transmembrane region, and an intracellular region. (B) Phylogenetic tree of the EqPD-L1 sequence in relation to other vertebrate species. The bootstrap consensus tree was inferred from 1000 replicates with the neighbor-joining method using the MEGA 7.0 software. The scale indicates the divergence time. The GenBank accession numbers of nucleotide sequences used in these analyses are listed in Supplementary Table 3.

### Interaction of EqPD-1 and EqPD-L1

We evaluated the cellular localization of EqPD-1-EGFP and EqPD-L1-EGFP proteins in the overexpressed COS-7 cell lines and found them to be localized on the cell surface (Fig 3A). We developed soluble recombinant EqPD-1-Ig and EqPD-L1-Ig in the Expi293 Expression System to analyze interactions of EqPD-1 and EqPD-L1 proteins. EqPD-1-Ig and EqPD-L1-Ig were successfully purified from culture supernatants and confirmed to be dimerized by disulfide bonds in the hinge region of rabbit IgG (Fig 3B).

**Figure 3.**
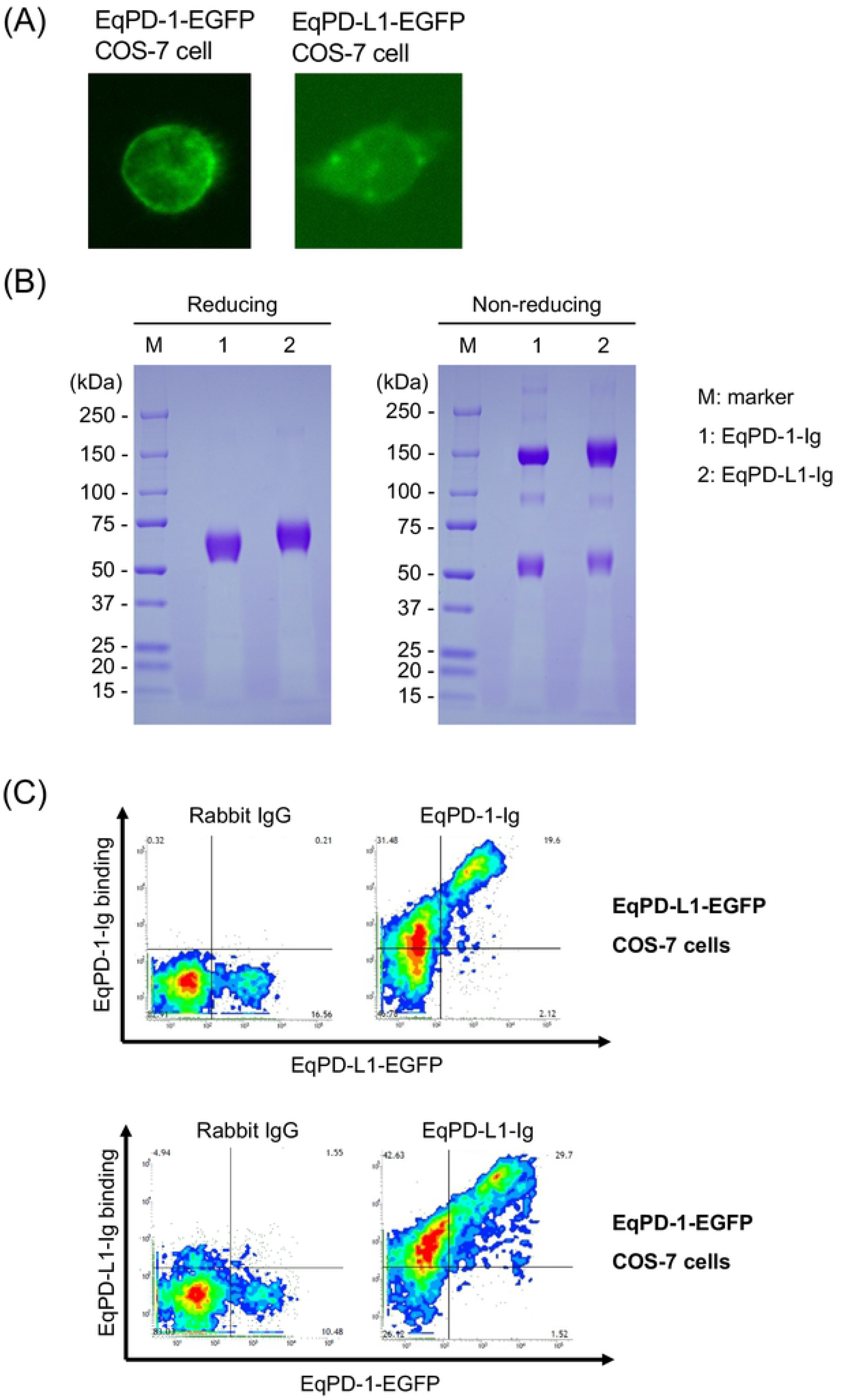
Establishment of EqPD-1- or EqPD-L1-expressing cells and Ig fusion soluble proteins. (A) EqPD-1-EGFP or EqPD-L1-EGFP-expressing COS-7 cell. The subcellular distributions of EqPD-1-EGFP and EqPD-L1-EGFP in transfected COS-7 cells were analyzed using a fluorescence microscope. (B) Production and purification of Ig fusion EqPD-1 and EqPD-L1 proteins. EqPD-1-Ig and EqPD-L1-Ig were purified from the culture supernatant and analyzed with SDS-PAGE. (C) Interaction of EqPD-1 and EqPD-L1. EqPD-1-EGFP or EqPD-L1-expressing COS-7 cells were incubated with EqPD-L1-Ig or EqPD-1-Ig, respectively. The binding of the Ig fusion proteins was labeled using Alexa Flour 647 conjugated anti-rabbit IgG antibody and analyzed by flow cytometry.

We used flow cytometry to analyze the interactions of EqPD-1-Ig or EqPD-L1-Ig with EqPD-L1-EGFP- or EqPD-1-EGFP-expressing cells, respectively. This revealed that EqPD-1-Ig binding to EqPD-L1-EGFP-expressing cells depends on the expression level of EqPD-1-EGFP (Fig 3C). Additionally, we confirmed that EqPD-L1-Ig binds to EqPD-1-EGFP-expressing cells in an expression dependent manner (Fig 3C).

### Cross-reactivity of anti-bovine PD-L1 mAbs against EqPD-L1

We evaluated cross reactivity of our previously established anti-bovine PD-L1 mAbs [5, 20] against EqPD-L1 and found that two out of the four tested mAbs (5A2-A1 and 6C11-3A11) detected EqPD-L1-EGFP overexpressed on COS-7 cells (Fig 4A). Of all the tested mAbs, 6C11-3A11 showed the strongest binding to EqPD-L1-EGFP-expressing cells (Fig 4A). We also tested the reactivity of 5A2-A1 and 6C11-3A11 mAbs against fresh and stimulated equine PBMCs and found that 6C11-3A11 mAb binds to both of fresh and stimulated PBMCs (Fig 4B and C). PD-L1 expression was upregulated on PBMCs through stimulation with PMA and ionomycin (Fig 4C).

**Figure 4.**
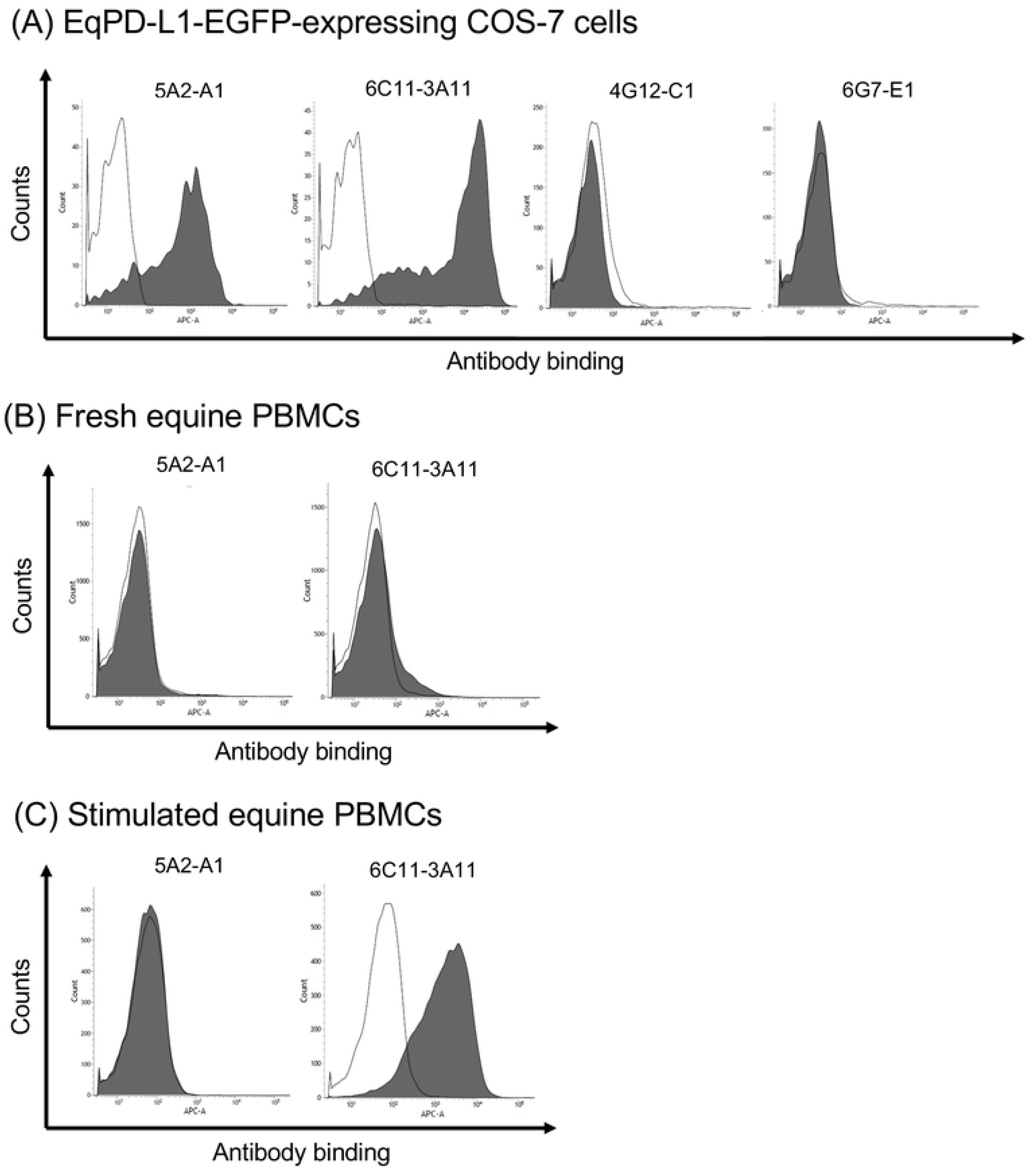
Cross-reactivity of anti-bovine PD-L1 mAbs against EqPD-L1. (A–C) Binding activities of anti-bovine PD-L1 mAbs (5A2-A1, 6C11-3A11, 4G12-C1, and 6G7-E1) to (A) EqPD-L1-EGFP-expressing COS-7 cells, (B) fresh equine PBMCs, and (C) equine PBMCs stimulated with PMA and ionomycin for 24 h. The binding of the primary mAbs was labeled with APC conjugated anti-rat Ig antibody and analyzed by flow cytometry. Rat IgG_1_ and IgG_2a_ controls were used as isotype-matched negative controls.

### Inhibition of EqPD-1/EqPD-L1 binding by anti-PD-L1 mAbs

We used ELISA to investigate whether the cross-reactive anti-bovine PD-L1 mAbs interfered with the interaction of EqPD-1/EqPD-L1. The 6C11-3A11 mAb, blocked the binding of EqPD-L1-Ig to EqPD-1-Ig in a dose-dependent manner, but the 5A2-A1 did not (Fig 5).

**Figure 5.**
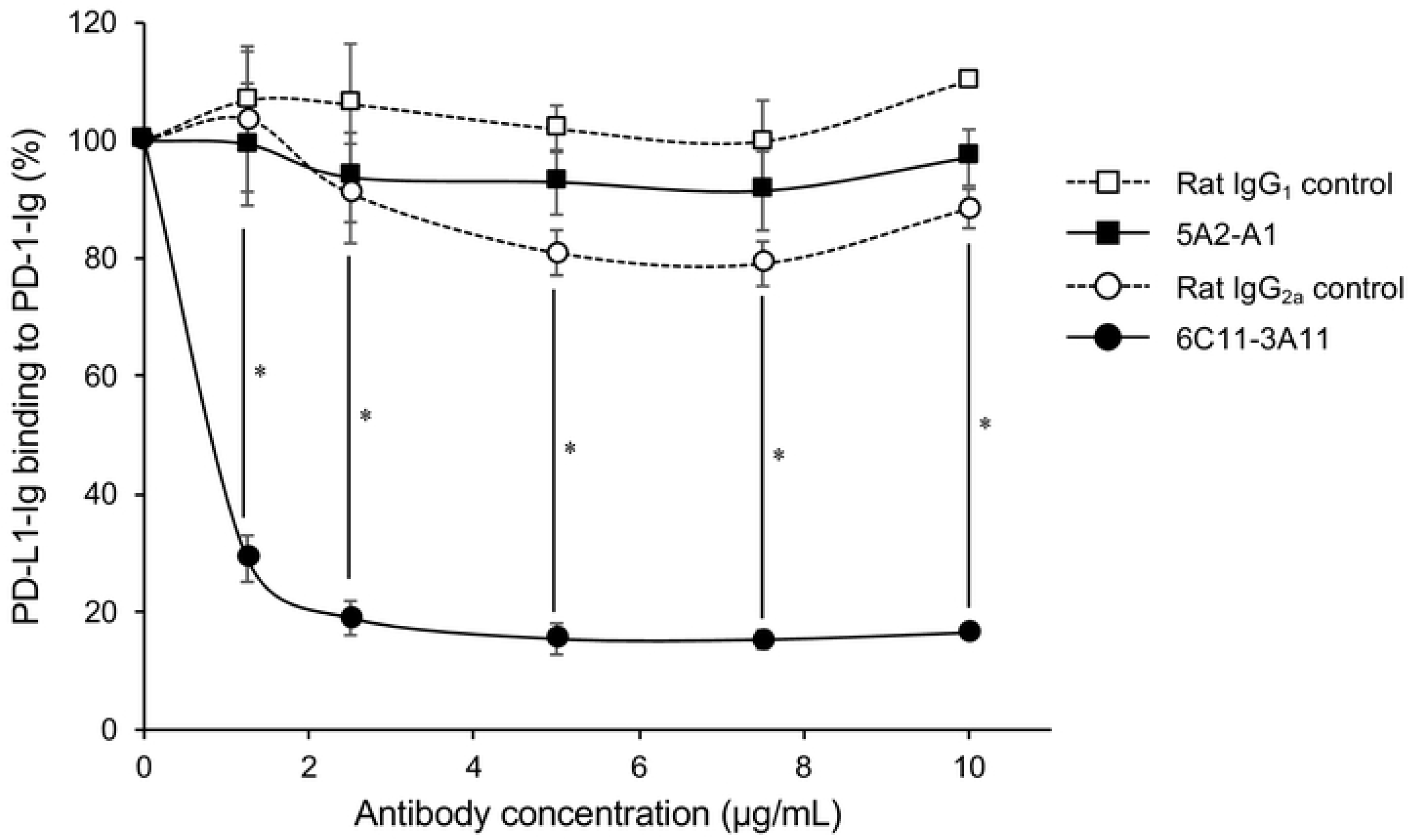
Inhibition of equine PD-1/PD-L1 binding by anti-PD-L1 mAbs. The blocking effect of anti-PD-L1 mAb on the binding of EqPD-L1-Ig to EqPD-1-Ig. EqPD-1-Ig was coated on a microwell plate. Biotinylated EqPD-L1-Ig was preincubated with various concentrations of anti-PD-L1 mAb (5A2-A1 or 6C11-3A11), and then incubated in the coated microwell plate. Rat IgG_1_ and IgG_2a_ controls were used as isotype-matched negative controls. Each curve represents the relative binding of EqPD-L1-Ig preincubated with antibodies compared to no-antibody control. Each point indicates the average value of three independent experiments. Significant differences between each treatment were identified using Tukey’s test. An asterisk (*) indicates *p* < 0.05.

### Immunohistochemical analysis of PD-L1 in EMM

PD-L1 has been shown to be upregulated on many types of tumors in dogs and humans [11, 12, 21]. Among canine malignant cancers, malignant melanoma has the highest positive rates for PD-L1 expression [11]. Gray horses are susceptible to melanoma and around 80% of them develop EMM in their lifetimes [3]. We hypothesized that PD-L1 plays a role in the development of EMM. Hence, we analyzed the expression of PD-L1 in tumor tissues of EMM by immunohistochemistry. PD-L1 was detected in all EMM samples (*n* = 4, Fig 6B).

**Figure 6.**
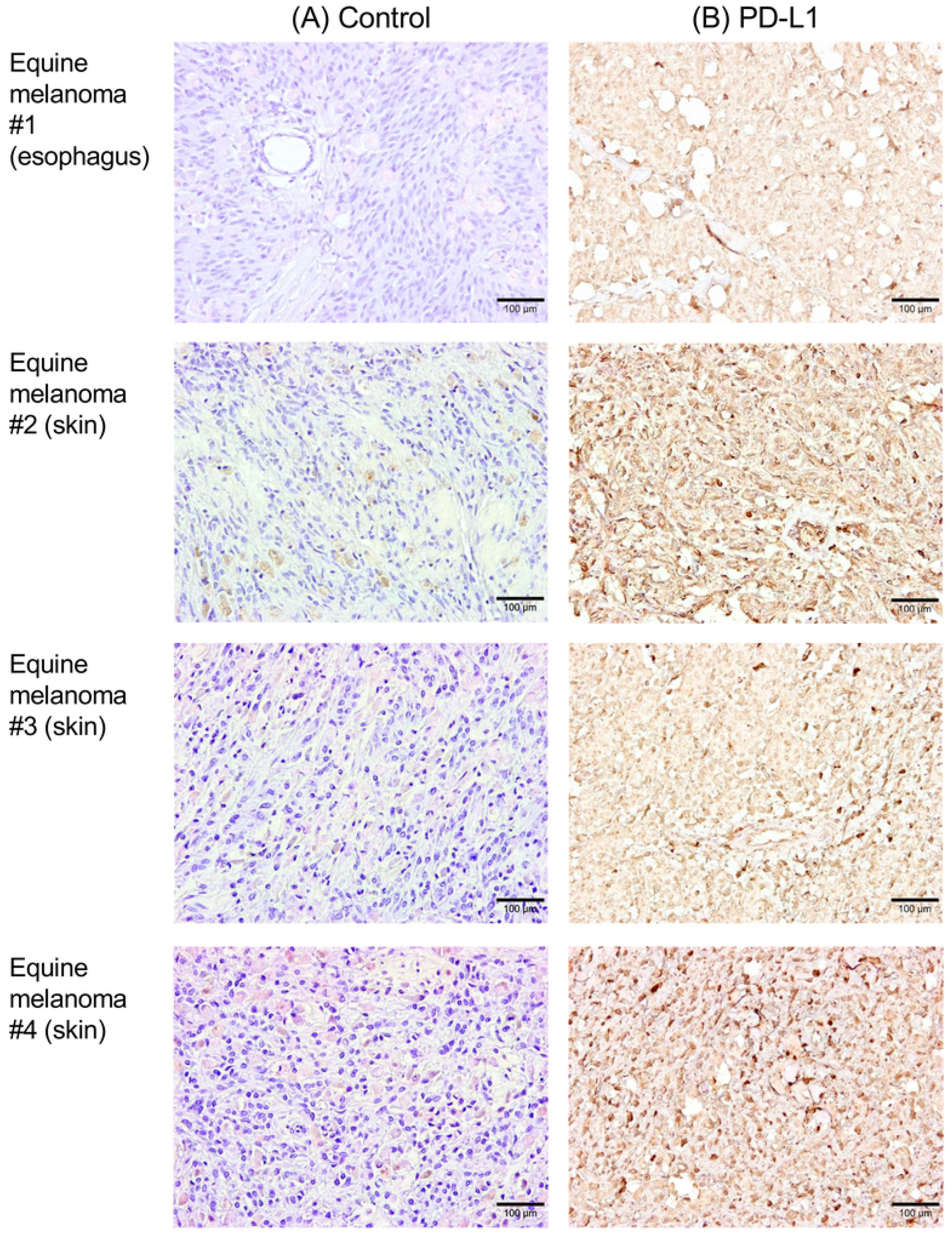
Immunohistochemical analysis of PD-L1 in EMM. Immunohistochemical staining of PD-L1 in tumor tissues of horses with melanoma (#1–#4). Each section was stained (A) without a primary antibody (control) or (B) using anti-bovine PD-L1 mAb (6C11-3A11). Further information of tumor specimens is shown in Supplementary Table 2.

### Immune activation in equine PBMCs by anti-PD-L1 mAb

We analyzed immune activation effects by PD-1/PD-L1 inhibition in the PBMC culture assays using anti-PD-L1 blocking mAb, 6C11-3A11. We found that PD-L1 blockade by the mAb 6C11-3A11 significantly induced IFN-γ production by equine PBMCs under stimulation with SEB (Fig 7A). Additionally, production of IL-2 was increased by PD-L1 inhibition (Fig 7B). These results indicate that PD-1/PD-L1 blockade enhanced Th1 cytokine production in equine PBMCs, suggesting that the anti-PD-L1 blocking antibody may have an application as an immunomodulatory agent for horses.

**Figure 7.**
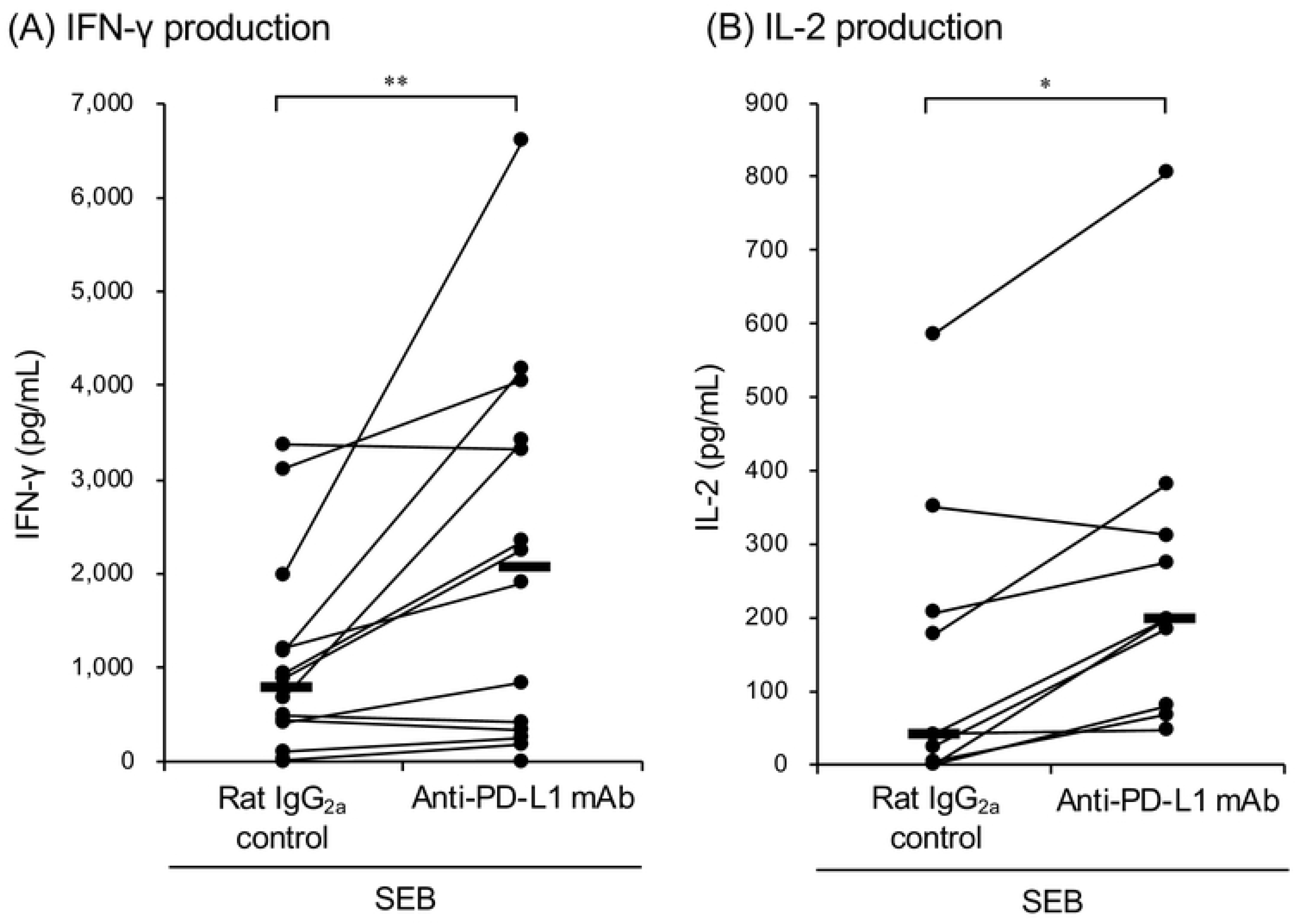
Effect of PD-L1 blockade on IFN-γ and IL-2 production. PBMCs isolated from healthy horses were cultured with anti-PD-L1 mAb (6C11-3A11) or rat IgG_2a_ control in the presence of SEB. The culture supernatants were harvested three days later and IFN-γ and IL-2 concentrations were measured by ELISA (IFN-γ: *n* =14 and IL-2: *n* = 9). Significant differences between each treatment were identified using Wilcoxon signed-rank test. Asterisks (* and **) indicate *p* < 0.05 and < 0.01, respectively.

## Discussion

The greater longevity of the horse population has increased the risks of chronic diseases, such as laminitis, pituitary pars intermedia dysfunction, recurrent airway obstruction, osteoarthritis, and neoplasia, and increased multimorbidity in horses [22, 23]. However, few treatments are available for chronic diseases in horses, including malignant tumors. Hence, new treatment options are being sought.

Malignant melanoma is one of the most common cutaneous neoplasia in horses [24] Surgical treatment is a successful in the early stages of disease, but it is not feasible in cases with multiple tumor burdens and metastases. Although a number of systemic treatments have been tested, no effective systemic therapy is currently available for EMM. To overcome the current situation, novel therapeutic strategies, including immunotherapy, are warranted for EMM.

A variety of immunotherapies have been developed and tested in clinical trials to treat tumors in humans, and immune checkpoint inhibitors such as anti-PD-1 and anti-PD-L1 antibodies are currently used with notable success for the treatment of multiple human cancers [25, 26]. Blockade therapy using anti-PD-L1 antibody resulted in long-term tumor regression and prolonged progression free survival in advanced melanoma in humans [26]. Based on these advancements in human medicine, immune checkpoint inhibitors may reasonably be expected to yield equally promising results in the treatment of EMM [4]. However, as yet no studies have been conducted on the PD-1/PD-L1 pathway in horses.

Our recent research revealed that the PD-1/PD-L1 pathway plays critical roles in immune exhaustion and disease progression in bovine chronic infections and canine malignant cancers [6–16]. Moreover, we established anti-PD-L1 and anti-PD-1 blocking antibodies for therapeutic application in cattle and dogs [13–15]. Clinical studies have confirmed the antiviral, antibacterial, and antitumoral effects of antibody treatments [13–17]. However, the blockade effect of the PD-1/PD-L1 pathway had not been tested in horses. Hence, we aimed to identify nucleotide sequences of EqPD-1 and EqPD-L1 and evaluate the function of our anti-bovine PD-L1 mAbs using *in vitro* assays.

We found that one of the anti-bovine PD-L1 mAbs (6C11-3A11) recognized EqPD-L1 strongly, blocked the interaction of EqPD-1/EqPD-L1 and enhanced the Th1 cytokine response *in vitro*. This anti-PD-L1 mAb may be used to aid investigation into the expression and immunological function of PD-L1 in future horse studies. Additionally, we discovered that PD-L1 is expressed in EMM tumor tissues. Further studies are required to analyze expression of PD-L1 in other horse tumors and chronic diseases.

The mechanism of PD-L1 upregulation during EMM progression has yet to be elucidated. Generally, PD-L1 expression is regulated by a substantial number of mediators including inflammatory cytokine signaling, oncogenic signaling, microRNAs, genetic alteration of the PD-L1 locus, and post-translational regulators [27]. In gray horses, a gene duplication in intron 6 of *STX17* (synataxin 17) contributes a *cis-acting* regulatory mutation resulting in a very high incidence of EMM [28]. This gene duplication induces constitutive activation of the extracellular signal-regulated kinase (ERK) pathway and melanomagensis in EMM [29, 30]. The MEK-ERK signaling pathway regulates PD-L1 gene expression via crosstalk with inflammatory cytokine signaling including the IFN-γ-STAT1 pathway [31–33]. Hence, the regulatory mechanism of PD-L1 expression in gray horses merits investigating as a natural model of tumorigenesis.

Our results indicate that the PD-1/PD-L1 pathway offers a potential target for immunotherapy against EMM. In future immunotherapy applications, blocking antibodies should be engineered into suitable forms for administration to horses. Chimeric antibodies, for instance, may facilitate clinical trial research into the clinical efficacy of anti-PD-L1 antibody in the treatment of EMM. Further research is required to develop this novel immunotherapy strategy in horses.

## Author Contributions

SK, TO, NM, SM, and KO: designed the work; GO, TO, YN, EM, and AK,: performed the experiments; RA, NS, DM, OI, YK, and YS: provided intellectual input, field samples, laboratory materials, reagents, and/or analytic tools; GO, SK, TO, and AK: acquired, analyzed, and interpreted the data; GO and SK: wrote the manuscript; SK, TO, NM, AK, NS, DM, OI, YK, YS, SM, KO: revised the manuscript; all authors: approved the final version of the manuscript.

## Acknowledgments

We are grateful to Dr. Hideyuki Takahashi, Dr. Yasuyuki Mori, and Dr. Tomio Ibayashi for valuable advice and discussions. We would like to thank Enago (www.enago.jp) for the English language review.

